# A novel antiviral strategy targeting human metapneumovirus through pH modulation in human airway epithelial cells

**DOI:** 10.64898/2026.02.11.704945

**Authors:** Ivana A. Daniels, Benjamin Gaston, Jessica Saunders, Laura Smith, Taiya Edwards, Nastaha Tilston, Ryan F. Relich, Andrew Lunel, Michael D. Davis

## Abstract

Human metapneumovirus (hMPV) is a major cause of respiratory infections, particularly among infants, older adults, and immunocompromised individuals, yet no approved vaccines or targeted antiviral therapies are currently available. pH-regulated processes, including airway epithelial physiology, endosomal acidification, and viral fusion mediated by the fusion (F) protein, are critical for hMPV infection. This study evaluates PHOH-001, an inhaled alkaline buffer, as a potential therapeutic strategy to modulate airway epithelial pH and inhibit hMPV infection. Using a recombinant hMPV expressing green fluorescent protein (rhMPV-GFP), viral replication was assessed in primary human airway epithelial cells (HAECs). PHOH-001 significantly reduced GFP expression at 72 hours post-infection in both submerged and air-liquid interface (ALI) cultures, with effects comparable to those of the endosomal acidification inhibitor bafilomycin A1. In Vero E6 cells, used as a mechanistic *in vitro* model, PHOH-001 increased extracellular and intracellular pH in a concentration-dependent manner and correspondingly reduced hMPV infection. In HAECs, PHOH-001 reduced viral replication, as measured by TCID_50_ assays of infectious virus, and inhibited syncytium formation, a key step in viral spread. Furthermore, PHOH-001 altered F protein localization and was associated with changes in actin organization, consistent with impaired viral spread. Collectively, these findings demonstrate that PHOH-001 alters multiple pH-dependent steps in hMPV infection *in vitro* and support airway pH modulation as a potential antiviral strategy against hMPV.

## Introduction

Human metapneumovirus (hMPV) is a leading cause of acute respiratory tract infections worldwide. Like other members of the family *Pneumoviridae*, hMPV possesses a non-segmented, negative-sense, single-stranded RNA genome, and viral particles are enveloped and adopt spherical morphologies (1, 2). Identified in 2001, hMPV has become recognized as a significant human respiratory pathogen. Disease manifestations can range from mild upper respiratory illness, including cough, nasal congestion, and fever, to severe lower respiratory disease such as bronchiolitis and pneumonia (3, 4). Severe disease occurs most frequently among infants, older adults, and immunocompromised individuals and is comparable in clinical severity to respiratory syncytial virus (RSV) infection (1, 5, 6). Nearly all children are infected by five years of age, and reinfections occur throughout life due to incomplete or short-lived immunity (7). Recent studies have shown that pediatric lower respiratory tract infections, many caused by hMPV, lead to a twofold increased risk of adult death from respiratory disease (8). Despite its high prevalence and significant contributions to long-term health outcomes, no hMPV-specific vaccines or therapies are approved for clinical use (9-11).

Like other pneumoviruses, hMPV infection begins with virion attachment to the respiratory epithelium, followed by fusion of the viral and host membranes, a process mediated exclusively by the viral fusion (F) protein (2). Unlike RSV, hMPV entry is more heterogeneous and can occur either by direct fusion at the plasma membrane or following endocytosis, with fusion occurring at endosomal membranes to release the viral genome into the cytoplasm (12-15). While direct fusion contributes to hMPV pathogenesis, multiple studies suggest that pH-dependent mechanisms, such as endocytosis and macropinocytosis, represent the predominant pathways for viral entry (13, 16, 17). These pathways are considered pH dependent, as endosomal maturation is accompanied by progressive acidification, which can promote conformational changes in the F protein that enable membrane fusion within intracellular compartments. Consequently, efficient endosomal trafficking and maturation are critical for successful genome release and downstream viral gene expression and replication. This requirement has implicated pH as an important regulator of F-protein activation and has fueled ongoing debate regarding the role of pH in hMPV entry.

Early biochemical and mutational analyses indicated that certain hMPV strains undergo robust low-pH-triggered conformational changes in the F protein, consistent with an endosomal fusion mechanism (18). However, subsequent studies demonstrated that pH-triggered fusion occurs only in a subset of strains and that many isolates fuse efficiently at neutral pH (19-21). To better understand the structural basis for this variability, analyses of prefusion-stabilized hMPV F reveal that the protein is stable at neutral pH and lacks the conserved pH-sensing residues typically required for acid-triggered fusion in other viral fusogens (22). Together, these observations support the view that pH sensitivity is strain dependent and likely influenced by determinants within the F protein heptad repeats, cleavage loop, and hydrophobic fusion peptide (20-22).

Despite these conflicting data regarding pH-triggered fusion, pH remains a biologically relevant antiviral target. hMPV primarily uses endocytic pathways such as clathrin-mediated endocytosis and macropinocytosis to access acidified endosomes, suggesting that endosomal pH may modulate entry efficiency (13, 23). Additionally, host airway surface liquid (ASL) pH regulates epithelial defense mechanisms, protease activity, and viral stability, and perturbations in ASL pH can influence infection dynamics (24-29). Lastly, F-protein conformational landscape remains sensitive to environmental factors, including ionic strength and protonation of specific residues, which may shift the balance between the prefusion and post-fusion states (22, 30-32). Together, these observations suggest that therapeutic modulation of pH, either by disrupting endosomal maturation or by altering the extracellular airway environment, may interfere with critical steps in viral entry. However, whether modulation of airway epithelial pH can be therapeutically leveraged to disrupt hMPV infection and F-protein-mediated viral spread in primary airway epithelium remains unclear.

Based on this rationale, we hypothesized that pharmacological elevation of airway epithelial cell pH would impair hMPV infection by disrupting pH-sensitive entry and post-entry processes. To test this hypothesis, we developed PHOH-001 (formerly “Optate”, Atelerix Life Sciences, Indianapolis, IN), an inhaled alkaline glycine buffer that has been shown to safely raise human airway pH *in vivo* (33, 34). In primary human airway epithelial cells (HAECs) *in vitro*, PHOH-001 raises intracellular pH and has been associated with altered expression of genes involved in endosomal trafficking, including *Rab5*, *Rab*7, *Rab9*, and *Rab11* (33). PHOH-001 also exhibits broad antiviral activity, reducing SARS-CoV-2 and RSV infection in HAECs (33, 35). Here, we examine the effects of PHOH-001 on hMPV entry, replication, and F-protein-mediated membrane fusion in organotypic human airway epithelial cultures to assess the potential of pH modulation as a therapeutic strategy for hMPV infection.

## Materials and Methods

### Cells and virus

Primary HAECs from the lower respiratory tract (bronchial) were obtained from donors at Indiana University School of Medicine. Pediatric primary HAECs were purchased from Lifeline Cell Technology (#07783) (Supplemental Table 1). Cells were cultured under submerged or air-liquid interface (ALI) conditions for 7 or 21 days, respectively, as previously described (33). Experiments with HAECs were conducted at passages 2-6. Vero E6 cells (African green monkey kidney cells) were maintained in Dulbecco’s Modified Eagle Medium (DMEM; Gibco, #11965092) supplemented with 10% non-heat-inactivated fetal bovine serum (FBS; Cytiva, SH3008003) and 1% penicillin/streptomycin (P/S; Gibco, #15140122) at passage 4. A green fluorescent protein-expressing recombinant human metapneumovirus (rhMPV-GFP; ViraTree, MPV-GFP1, #M121, CAN97-83 [A2]) was used at virus passages 4-5. Viral stocks were propagated in LLC-MK2 cells (ATCC CCL-7) and stored in suspension medium consisting of Opti-MEM, sucrose, and L-glutamic acid (36). Viral titers were ≥7.0 log_10_ TCID_50_ mL^-1^.

All experiments were carried out in humidified conditions at 37°C under 5% CO_2_.

### Viral growth kinetics

Vero E6 cells and HAECs were seeded in black-walled, clear-bottom 96-well plates (Corning, #3603) and grown to confluency. Vero E6 and HAEC monolayers were rinsed with phosphate-buffered saline (PBS; Gibco, #10010023) and infected with rhMPV-GFP at a multiplicity of infection (MOI) of 0.01, 0.1, or 1 (n=3). The virus was diluted in FluoroBrite™ DMEM (Gibco, A1896701), a phenol red-free medium, to minimize autofluorescence during live cell imaging. GFP signals were quantified at specified timepoints using a FLUOstar Omega plate reader (BMG Labtech), and viral growth kinetics were plotted as relative light units (RLU). Supernatants collected at each timepoint were titrated using TCID_50_ assays to determine infectious viral titers.

### TCID_50_ assay

Vero E6 cells were seeded at 1.2x10^4^ cells per well in 96-well plates (Corning, #3360). After 24 h, monolayers were inoculated with 10-fold serial dilutions of infected supernatant. At 5-7 days post-infection (dpi), wells were scored for the presence of GFP-positive infection. Viral titers were calculated and expressed as TCID_50_ mL^-1^. The limit of detection (LOD) for the TCID_50_ assay was calculated based on the lowest dilution tested (10^-1^) and an inoculation volume of 200 µL per well, resulting in an LOD of 1.7 log_10_ TCID_50_/mL.

### GFP-based quantification of infection via fluorescence microscopy

Confluent HAEC monolayers in 12-well plates were infected with rhMPV-GFP (MOI=1). PHOH-001 was prepared by a compounding pharmacy at 120 mM and verified for purity, potency, osmolality (∼330 mOsm), pH (9.8), and sterility (IND #139144; Arena District Pharmacy) (34). PHOH-001 (120 mM) or control (PBS; pH 7.2) was co-administered with rhMPV-GFP diluted in Opti-MEM (Gibco, #31985070) (MOI=1) (1:1 dilution) for 4 h; Bafilomycin (100 nM; Millipore Sigma, #508409) served as an additional treatment group in ALI experiments and was co-administered with rhMPV-GFP for 4 h. Supernatants were collected at 4 h post-infection (hpi) and every 24 h for 72 h for both submerged and ALI cultures. Submerged cultures were treated daily during medium changes; ALI cultures were treated apically for 20 min twice daily with daily basolateral medium changes supplemented with 0.3 µg mL^-1^ TPCK-treated trypsin (Thermo Scientific, #20233).

Images were acquired every 24 h for 3 days using an EVOS M5000 microscope (Thermo Fisher Scientific) with an EVOS™ Light Cube, GFP 2.0 (Thermo Fisher Scientific, AMEP4951) at 4x magnification (n=6). GFP-positive regions were segmented in Fiji (37) using noise-reduction parameters and thresholding, followed by particle analysis. The GFP-positive area was quantified and compared across treatment groups to assess infection at 72 hpi.

### Automated Western blot

At 72 hpi, HAECs from infected and treated submerged and ALI cultures were lysed in 50 µL RIPA buffer (Sigma Aldrich, R0278) supplemented with 1% protease inhibitor (PI) (Millipore, #539134). Lysates were centrifuged at 14,000 x g for 10 min at 4°C, and supernatants were stored at -80°C. Protein concentration was measured using the Pierce BCA Protein Assay Kit (Thermo Fisher Scientific, #23250). Samples were diluted in 1X Sample Buffer to 0.25 µg µL^-1^ and analyzed using the Jess automated western capillary system (ProteinSimple, #004-650). GFP was detected using a rabbit monoclonal antibody (1:50; Cell Signaling Technology; 2956) normalized to β-actin detected with a mouse monoclonal antibody (1:25; Cell Signaling Technology; 3700). hMPV F protein was detected using a rabbit polyclonal antibody (1:25; AntibodySystem, PVV22901) normalized to β-actin (1:25; Cell Signaling Technology; 3700). Chemiluminescent signals were quantified using Compass for SW software (ProteinSimple).

### GFP enzyme-linked immunosorbent assay (ELISA)

HAECs from infected and treated submerged and ALI cultures were lysed at 72 hpi in 50 µL 1X Cell Extraction Buffer PTR (CatchPoint® SimpleStep GFP ELISA; Abcam, ab229403). Protein concentrations were measured (Pierce BCA Protein Assay Kit; Thermo Fisher Scientific), and lysates were centrifuged at 14,000 x g for 10 min at 4°C and stored at -80°C. Prior to analysis, samples were diluted to the required concentration in 1X Cell Extraction Buffer PTR, and a standard curve was generated using recombinant GFP provided with the kit. HRP fluorescence was quantified using a FLUOstar Omega plate reader following the manufacturer’s protocol, and GFP concentrations were calculated using a four-parameter logistic regression model.

### Dose-response assays

Vero E6 cells were seeded at 1.2x10⁴ cells per well in 96-well plates. Upon confluency, cells were infected with rhMPV-GFP (MOI=0.1) and immediately treated with PHOH-001 at 120 mM, followed by 1:2 serial dilutions across all 12 columns (n=8). Plates were incubated with PHOH-001 for 48 h. Fluorescence was measured using a FLUOstar Omega plate reader and whole-plate images were acquired using an Odyssey M Imager (LI-COR). To ensure accurate interpretation of the results, the pH of the PHOH-001 stock and its serial dilutions were assessed simultaneously with the experiment using a pH meter (Mettler-Toledo).

### Intracellular pH assay

Vero E6 cells were seeded in black-walled, clear-bottom 96-well plates. Once confluent, cells were washed with Live Cell Imaging Solution (Thermo Fisher Scientific, A59688DJ) and stained with pHrodo™ Red Intracellular pH Dye according to the manufacturer’s instructions (Thermo Fisher Scientific, P36600). Cells were treated with PHOH-001 at 120 mM followed by 1:2 serial dilutions in FluoroBrite™ DMEM across 12 columns (n=8). pHrodo™ Red fluorescence was quantified using a FLUOstar Omega plate reader (excitation 544 nm, emission 590 nm) at 30 min post-treatment. Immediately following fluorescence measurements, plates were imaged using an Odyssey M full-plate imager to ensure consistency between quantitative and imaging-based readouts. Higher-resolution fluorescence images were subsequently acquired using an EVOS M5000 microscope (Thermo Fisher Scientific) equipped with an EVOS™ Light Cube, RFP 2.0 (Thermo Fisher Scientific, AMEP4952) at 4x magnification.

### Syncytium assay

Confluent HAEC monolayers in black-walled, clear-bottom 96-well plates were co-inoculated with rhMPV-GFP (MOI=1) and PHOH-001 or PBS (1:1 dilution) for 4 h. At 4 hpi, the inoculum was removed, the monolayers were rinsed with PBS, and fresh medium containing PHOH-001 or PBS (1:1 dilution) was added. Cultures were maintained for 72 h with daily medium replacement containing the corresponding treatment.

At 72 hpi, monolayers were fixed with 4% paraformaldehyde (PFA) for 15 min and permeabilized with PBS containing 0.1% Triton X-100 (Sigma Aldrich, #9036-19-5) for 20 min. GFP signal remained stable after fixation and did not require additional staining. Upon fixation, the cell monolayers were blocked for 1 h using 5% donkey serum (Rockland, D220-00-0050). Cell membranes were then stained with a monoclonal antibody against E-Cadherin (1:200; BD Biosciences, #610181) and nuclei were stained with Hoechst 33342, Trihydrochloride, Trihydrate (Thermo Fisher Scientific, H1399) following the manufacturer’s instructions. After washing, samples were incubated with goat anti-Mouse IgG 647 (Invitrogen, A32728) followed by a final wash step. Immunofluorescence imaging was acquired at 60x magnification using an EVOS M5000 microscope (Thermo Fisher Scientific) (n=25).

Syncytia were quantified using CellProfiler (v4.2.8) with a custom pipeline for segmenting nuclei, GFP-positive regions, and E-cadherin-defined cell borders. Nuclei within GFP-positive, membrane-bounded regions were counted, and cells containing ≥2 nuclei were classified as syncytia. The frequency of syncytia was calculated by dividing the number of nuclei within syncytia by the total number of nuclei per field and multiplying by 100, providing the percent of cells participating in syncytia for each treatment group. Total nuclei were used rather than total GFP-positive cells because imaging at 60x provided a limited field of view, and using total nuclei gave a consistent measure across images.

### Fusion protein analysis via confocal microscopy

HAECs were seeded on glass coverslips at 5.0x10^4^ cells per coverslip. Upon reaching confluency, cells were co-inoculated with rhMPV-GFP (MOI=1) and PHOH-001 or PBS (120 mM) (1:1 dilution) for 4 h. At 4 hpi, the inoculum was removed, the monolayers were rinsed with PBS, and fresh medium containing the corresponding treatment was added. Cultures were maintained for 72 h with daily medium and treatment replacement.

At 72 hpi, coverslips were fixed with 4% PFA for 15 min and permeabilized with PBS containing 0.1% Triton X-100 for 20 min. GFP signal remained stable after fixation and required no additional staining. Coverslips were blocked with 5% donkey serum for 1 h, then incubated with a monoclonal antibody against hMPV F protein (1:300; Abcam, ab94800) and phalloidin (Thermo Fisher Scientific, A22283) to label filamentous actin for cell segmentation. After washing, samples were incubated with goat anti-mouse IgG Alexa Fluor 647, followed by final washes and mounting with ProLong™ Gold antifade reagent containing DAPI (Invitrogen, P36931).

Images were acquired on a Zeiss LSM 800 AxioObserver with Airyscan using a 20x and 63x Plan-Apochromat objective (n=25 cells per condition). Image analysis for 63x images was performed using CellProfiler. Phalloidin staining was used to delineate individual cell boundaries and enable segmentation of single cells, while GFP signal was used to identify infected cells. A plasma membrane-proximal peripheral region was generated by uniformly eroding the phalloidin-defined whole-cell mask inward by a fixed distance of 5 μm, and the cytoplasmic region was defined as the remaining intracellular area. Integrated fluorescence intensity of F protein was measured within the whole-cell and peripheral regions by summing pixel intensity values within each region of interest. Peripheral enrichment was calculated on a per-cell basis as the ratio of integrated F protein intensity within the peripheral region to the integrated intensity within the whole cell and is reported as a percentage. Values were compared across treatment conditions.

### RT-qPCR

At 72 hpi, HAECs from infected submerged models treated with PHOH-001 or PBS were lysed, and total RNA was extracted using a RNeasy® Plus Mini Kit (Qiagen, #74134) following the manufacturer’s instructions. RNA samples were reverse transcribed to cDNA using the Maxima H Minus cDNA Synthesis Kit (Thermo Fisher Scientific, M1681) according to the manufacturer’s protocol. cDNA was eluted in 50 µL of Ambion Nuclease-Free Water (Invitrogen), quantified using a Biospectrometer (Eppendorf), and diluted to a final concentration of 100 ng per sample.

Primers, probe, and a synthetic gene block were designed based on the hMPV F protein gene sequence (NCBI Accession: AY145296, prototype A2 subtype CAN97-83). Primer and probe sequences were checked for cross-reactivity, self-dimerization, hairpin formation, and secondary structure using NCBI BLAST, OligoCalc, and OligoAnalyzer. Complete sequences are provided in Supplemental Table 2; primers and gene block were synthesized by Integrated DNA Technologies (IDT), and the probe was purchased from Applied Biosystems.

RT-qPCR was performed using a QuantStudio 3 instrument (Applied Biosystems, A28567) with TaqMan Fast Virus 1-Step Multiplex Master Mix (Applied Biosystems, 5555532). Absolute quantification was determined using a standard curve generated using tenfold serial dilutions of the gene block, each tested in triplicate (38). Each 25 µL PCR reaction contained 400 nM forward primer, 400 nM reverse primer, 250 nM TaqMan probe (FAM), 2 µL of cDNA, and TaqMan Fast Virus 1-Step MM; primer efficiencies may be found in supplemental data with corresponding experiments. Thermal cycling was performed using a fast run protocol: 5 minutes at 50 °C for reverse transcription, followed by 20 seconds at 95 °C for initial denaturation, and then 40 cycles of 3 seconds at 95 °C for denaturation and 30 seconds at 60 °C for annealing.

### Data analysis

All statistical analyses were performed using GraphPad Prism (version 10.6.1). For comparisons between two groups, a two-sample, two-tailed t-test was applied to data with a Gaussian distribution, whereas the Wilcoxon rank-sum test was used for non-Gaussian-distributed data. For experiments involving more than two groups, robust one-way ANOVA models were applied, followed by pairwise comparisons using Tukey’s post hoc test. For ELISA assays, a four-parameter logistic regression model was used according to the manufacturer’s instructions. Schematics were created using BioRender.com. Biological replicates represent independent donor-derived HAEC cultures, whereas technical replicates represent independent wells or fields within the same culture. To differentiate donor origins, biological replicates are color-coded accordingly. For all data displayed, circles represent PBS-treated controls, squares represent PHOH-001-treated samples, diamonds represent bafilomycin-treated samples, and triangles represent uninfected controls. Statistical significance was defined as a p-value ≤ 0.05.

### Imaging quantification codes

The CellProfiler pipelines and Fiji scripts used for imaging-based quantification of syncytium formation and F protein localization are available from the corresponding author upon request (A.L.).

### Bioethics and biosafety statement

All experiments involving primary HAECs were approved by the Indiana University Institutional Review Board (protocol #1408855616) and Institutional Biosafety Committee (#1127). Written informed consent was obtained from all donors. All procedures adhered to institutional and national guidelines for human subject research.

All experiments involving rhMPV-GFP were conducted in a biosafety level 2 (BSL-2) laboratory under appropriate containment conditions. Personnel were trained in handling infectious agents, and all waste was decontaminated according to institutional biosafety guidelines.

## Results

### Recombinant hMPV-GFP exhibits robust replication kinetics and validates GFP as a quantitative reporter in primary HAECs

To establish a rapid and quantitative approach for monitoring hMPV infection, we assessed the replication dynamics of a recombinant GFP-expressing hMPV (rhMPV-GFP; CAN97-83 [A2]) in Vero E6 cells and primary HAECs. Assay parameters and infection conditions were first optimized in Vero E6 cells prior to evaluation in HAECs (Supplemental Figure 1). HAEC cultures were then inoculated at multiplicities of infection (MOI) of 0.01, 0.1, and 1. GFP expression increased in a clear MOI- and time-dependent manner, with GFP signal expanding across the epithelial surface over the course of infection (Figure 1A). Quantification of infection by both relative light units (RLU) and infectious titers (TCID_50_) demonstrated that GFP signal closely paralleled viral replication kinetics across all inoculation parameters tested (Figure 1B). RLU values rose progressively at each time point and strongly correlated with corresponding increases in infectious titers, confirming that GFP signal provides a reliable and quantitative surrogate for productive hMPV replication in primary HAECs. These findings validate the rhMPV-GFP infection model as an effective tool for downstream mechanistic and antiviral studies in the airway epithelium.

**Figure 1.**
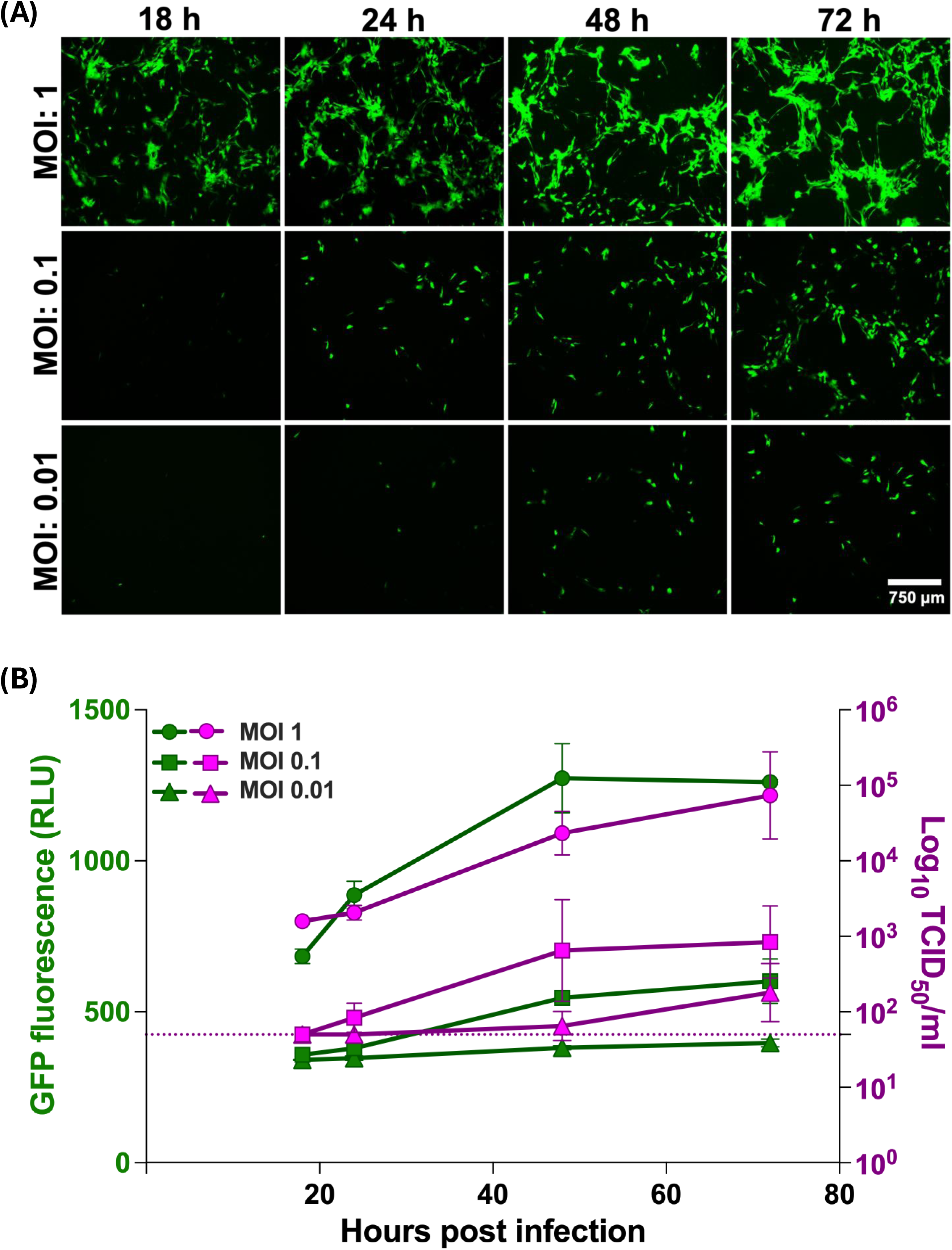
Validation of GFP fluorescence as a quantitative reporter of hMPV infection in primary HAECs. **(A)** Representative fluorescence images of HAECs infected with rhMPV-GFP at MOIs of 0.01, 0.1, or 1 at 18, 24, 48, and 72 hpi. GFP-positive signal is detectable by 24 hpi and increases over time. Images were acquired at 4x magnification. Scale bar=750 µm**. (B)** Quantification of GFP fluorescence (RLU; left y-axis) and infectious titers (TCID_50_/mL; right y-axis) over time. RLU and TCID_50_ increased in parallel in a MOI- and time-dependent manner. Dashed line indicates the TCID_50_ assay LOD (1.7 log10 TCID_50_/mL), with values below the limit plotted at the threshold. Data represents mean ± SD from three technical replicates.

### PHOH-001 significantly reduces hMPV infection in primary HAECs under submerged culture

To evaluate whether PHOH-001, an alkaline buffer, inhibits hMPV infection via pH modulation, primary HAECs were co-inoculated with rhMPV-GFP (MOI=1) and either PHOH-001 or PBS control buffer. Cells were treated daily with PHOH-001 or PBS post-inoculation (Figure 2A). Fluorescence microscopy at 72 hpi revealed reduced GFP fluorescence in PHOH-001-treated cells compared to controls (Figure 2B). Quantification using Fiji software confirmed a significant decrease in GFP-positive areas in PHOH-001-treated HAECs (Figure 2C). Western blot analysis confirmed decreased GFP expression in PHOH-001-treated cells (Figure 2D), and quantification of GFP normalized to β-actin confirmed a consistent reduction (Figure 2E). Additionally, ELISA assays were performed as a secondary protein readout, further supporting the western blot data by showing a significant reduction in GFP levels in PHOH-001-treated cells (Figure 2F). Together, these results demonstrate that PHOH-001 effectively inhibits hMPV infection in primary HAECs *in vitro*.

**Figure 2.**
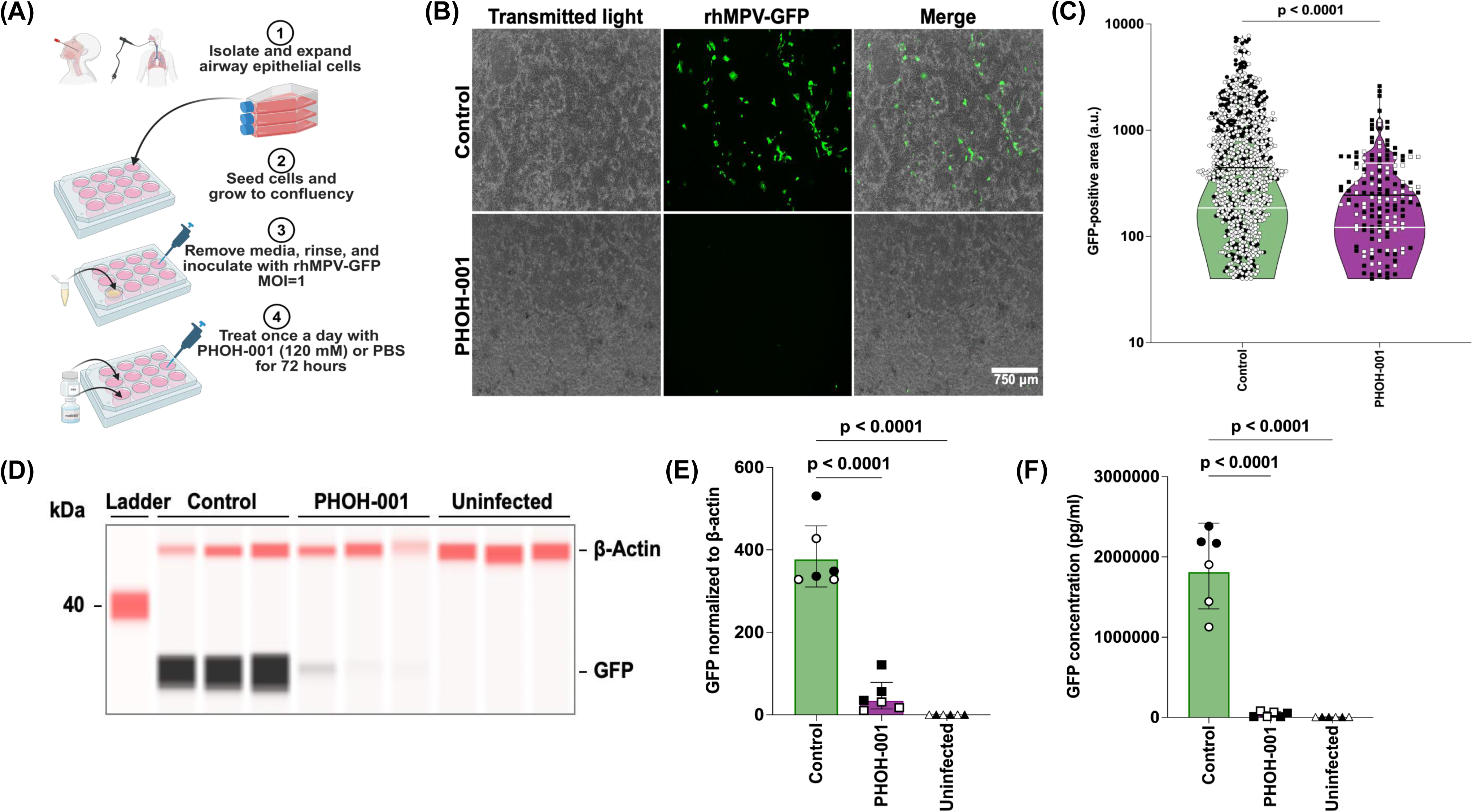
PHOH-001 reduces hMPV infection in submerged HAECs. **(A)** Experimental design. Primary HAECs were infected with rhMPV-GFP (MOI=1) and treated daily with PHOH-001 or PBS for 72 h. **(B)** Representative fluorescence images at 72 hpi show reduced GFP signal in PHOH-001-treated cells relative to PBS controls. Images were acquired at 4x magnification. Scale bar=750 µm. **(C)** Violin plot quantification of GFP-positive areas demonstrates GFP signal was significantly reduced in PHOH-001-treated cells compared to PBS. The violin shape is visually truncated to enhance readability; all individual data points are shown; black line=median, white lines=upper and lower quartiles. Data represents individual data points with mean ± SD from two biological replicates, each with six technical replicates (p<0.05, two-tailed unpaired Student’s t-test). **(D)** Western blot analysis of GFP expression at 72 hpi. **(E)** Quantification of GFP normalized to β-actin confirms reduced GFP levels in PHOH-001-treated cells. Data represents mean ± SD from two biological replicates, each with three technical replicates (p<0.05, one-way ANOVA). **(F)** ELISA quantification of GFP signal confirms a reduction in PHOH-001-treated cells. Data represents mean ± SD from two biological replicates, each with three technical replicates (p<0.05, one-way ANOVA).

### PHOH-001 significantly reduces hMPV infection in organotypic, differentiated primary HAECs

To assess whether PHOH-001 inhibits hMPV infection through pH modulation, primary HAECs were cultured under air-liquid interface (ALI) conditions, generating a differentiated, pseudostratified, ciliated epithelium that mimics the *in vivo* airway epithelium. Bafilomycin A1, a well-established inhibitor of endosomal acidification and a known inhibitor of hMPV entry, was included as a positive control (13, 19). HAECs were co-inoculated with rhMPV-GFP (MOI=1) and PHOH-001, bafilomycin, or PBS control buffer on the apical side of the membrane (Figure 3A). TPCK-treated trypsin was added to the basolateral media to cleave the F0 precursor and promote viral replication (15, 20, 36). Fluorescence microscopy at 72 hpi revealed a significant reduction in GFP signal in PHOH-001-treated cells compared to PBS-treated controls (Figure 3B). Quantification of GFP fluorescence using Fiji showed a significant reduction in GFP signal in PHOH-001-treated HAECs relative to PBS-treated controls (Figure 3C), with no significant difference observed between PHOH-001 and bafilomycin treatments. Western blot analysis also demonstrated decreased GFP expression in PHOH-001-treated cells (Figure 3D), with quantification showing significant reduction in GFP normalized to β-actin (Figure 3E). Additionally, ELISA analysis of GFP expression further supported the inhibitory effect of PHOH-001, showing a significant reduction in GFP levels in PHOH-001-treated cells compared to PBS controls (Figure 3F). Notably, no significant differences in GFP expression were found between PHOH-001 and bafilomycin-treated HAECs. Taken together, these data demonstrate that PHOH-001 effectively reduces hMPV infection in primary HAECs cultured at ALI, highlighting its potential as a therapeutic candidate for further translational applications.

**Figure 3.**
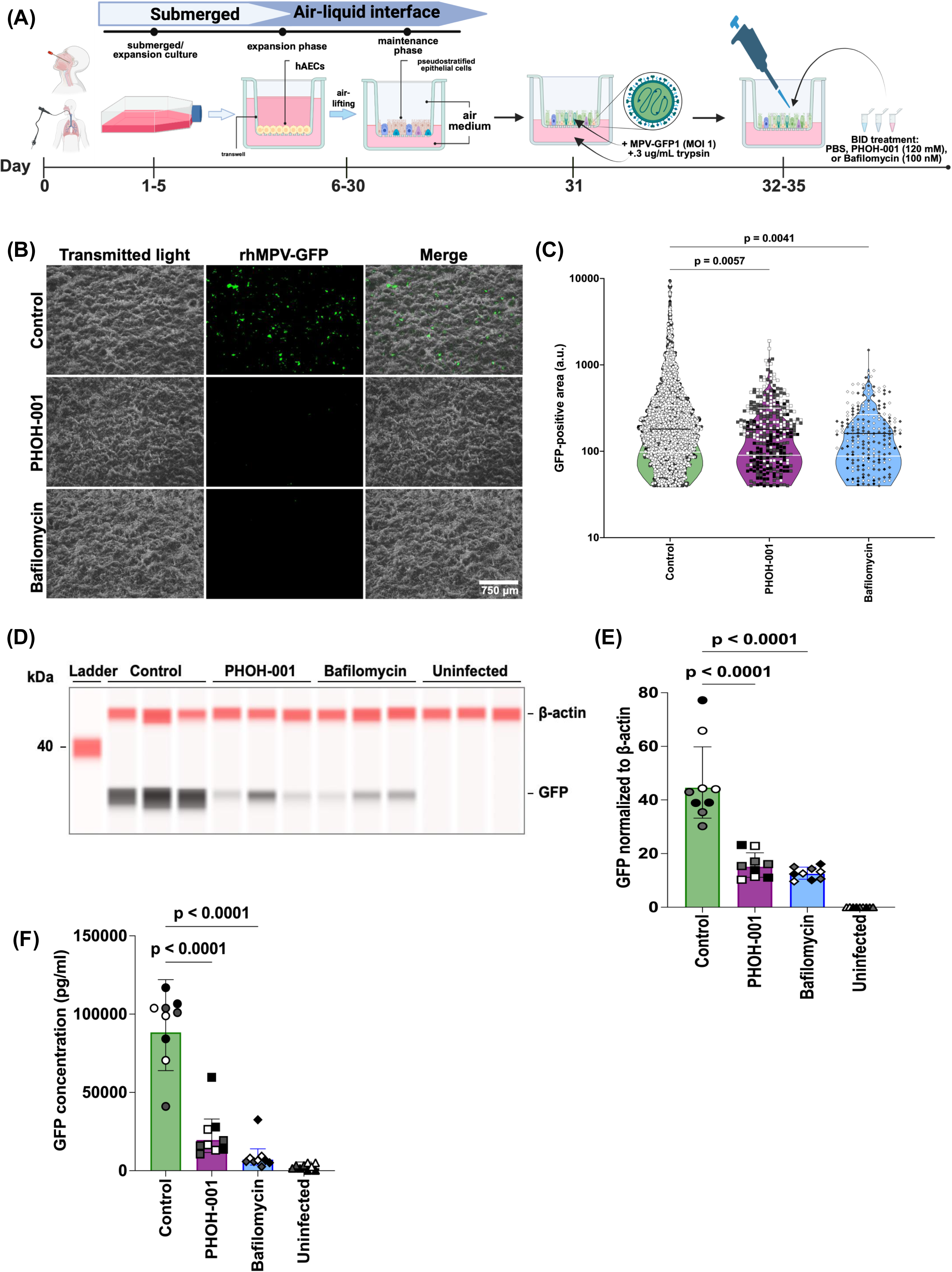
PHOH-001 reduces hMPV infection in HAECs cultured at ALI. **(A)** Experimental design; differentiated HAECs were inoculated with rhMPV-GFP (MOI=1) and treated apically with PHOH-001, bafilomycin, or PBS. Cells were treated twice daily, with once-daily basolateral media changes supplemented with TPCK-trypsin. **(B)** Representative fluorescence images at 72 hpi show reduced GFP signal in PHOH-001-treated cells relative to PBS controls. Images were acquired at 4x magnification. Scale bar=750 µm. **(C)** Violin plot quantification of GFP-positive areas demonstrates significant reduction in PHOH-001-treated cells compared to PBS, with no significant difference relative to bafilomycin. The violin shape is visually truncated to enhance readability; all individual data points are shown; black line=median, white lines=upper and lower quartiles. Data represents individual data points with mean ± SD from three biological replicates, each with six technical replicates (p<0.05, one-way ANOVA). **(D)** Western blot analysis of GFP expression at 72 hpi. **(E)** Quantification of GFP normalized to β-actin confirms reduced GFP expression in PHOH-001-treated cells relative to PBS, with no significant difference relative to bafilomycin. Data represents mean ± SD from three biological replicates, each with three technical replicates (p<0.05, one-way ANOVA). **(F)** ELISA analysis corroborates reduced GFP expression in PHOH-001-treated cells. Data represents mean ± SD from three biological replicates, each with three technical replicates (p<0.05, one-way ANOVA).

### PHOH-001 reduces hMPV infection in a dose-dependent manner correlated with pH modulation

To investigate the antiviral effects of PHOH-001 against hMPV, Vero E6 cells were infected with rhMPV-GFP (MOI=0.1) and treated with increasing concentrations of PHOH-001. Fluorescence imaging at 48 hpi revealed a concentration-dependent reduction in GFP signal (Figure 4A). Quantification of GFP fluorescence (RLU) and comparison to the corresponding PHOH-001 buffer pH showed that as GFP signal decreased, PHOH-001 pH increased (Figure 4B), suggesting that inhibition of hMPV infection is linked to pH modulation. Intracellular pH was assayed using the pH-sensitive dye pHrodo™ Red. Fluorescence measurements indicated that intracellular pH increased with higher PHOH-001 concentrations (Figure 4C-D; Supplemental Figure 2). Western blot and RT-qPCR analysis across a narrower dose range further confirmed this trend, showing reduced GFP protein and hMPV F gene expression with increasing PHOH-001 concentrations (Supplemental Figure 3). Together, these results demonstrate that PHOH-001 exerts a dose-dependent inhibitory effect on hMPV infection, which correlates closely with its ability to modulate extracellular and intracellular pH.

**Figure 4.**
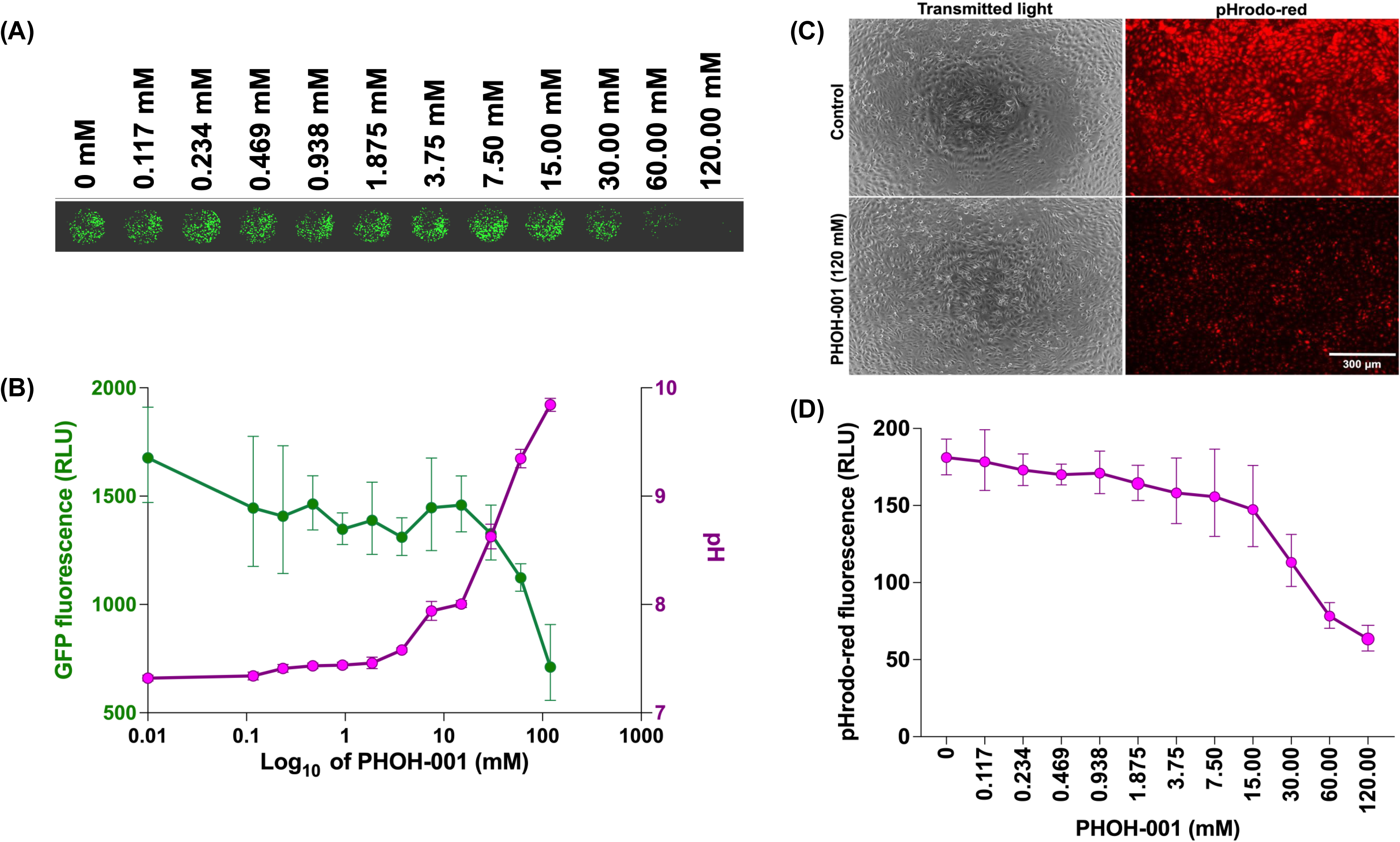
Dose-dependent effects of PHOH-001 on hMPV infection and pH. **(A)** Vero E6 cells were infected with rhMPV-GFP (MOI=0.1) and treated with increasing concentrations of PHOH-001. Whole-plate fluorescence images were acquired, showing a concentration-dependent reduction in GFP signal. The image shown corresponds to a single row of the 96-well plate. **(B)** Quantification of GFP signal (RLU; left y-axis) plotted alongside the pH of PHOH-001 buffer (pH; right y-axis). GFP signal decreased as PHOH-001 concentration increased, with a corresponding increase in pH. Data represents mean ± SD from eight technical replicates for RLU and three technical replicates for pH. **(C)** Representative fluorescence images of pHrodo™ Red intracellular pH assays in Vero cells treated with 0 mM or 120 mM PHOH-001. Images were acquired at 10x magnification. Scale bar=300 µm. **(D)** Quantification of pHrodo™ Red fluorescence in Vero E6 cells treated with increasing concentrations of PHOH-001. Fluorescence was measured by plate reader and expressed as RLU. Because pHrodo™ Red fluorescence increases with decreasing intracellular pH, the observed dose-dependent decrease in fluorescence indicates an increase in intracellular pH with higher PHOH-001 concentrations. Data represents mean ± SD from eight technical replicates.

### PHOH-001 reduces hMPV replication and syncytium formation in primary HAECs, a key step in viral spread

To further define the antiviral effects of PHOH-001 on hMPV infection, we examined viral replication kinetics and syncytium formation, a characteristic feature of hMPV-mediated spread between infected cells (17). We first assessed viral titers using a TCID_50_ assay, which demonstrated a significant reduction in infectious viral titers in the supernatants of PHOH-001-treated HAECs compared to controls (Figure 5A). This reduction in viral titers indicates that PHOH-001 effectively inhibits viral replication, a crucial step for hMPV to continue its infection cycle. Next, we examined syncytium formation, a hallmark of hMPV infection, which is critical for viral dissemination within the host. Syncytia are multinucleated giant cells formed when the virus induces fusion between infected cells and adjacent uninfected cells (15, 39). Fluorescence microscopy images showed that syncytia were notably reduced in PHOH-001-treated HAECs compared to PBS-treated controls, providing further evidence that PHOH-001 limits viral spread by impairing both replication and syncytium formation (Figure 5B). Quantification of syncytium formation in PHOH-001-treated and control cells revealed a significant reduction in syncytia in the treated samples, indicating that PHOH-001 potentially interferes with viral fusion and the propagation of infection (Figure 5C). Taken together, these findings indicate that PHOH-001 does not fully prevent hMPV entry but substantially impairs productive infection, as reflected by reduced viral replication and syncytium formation.

**Figure 5.**
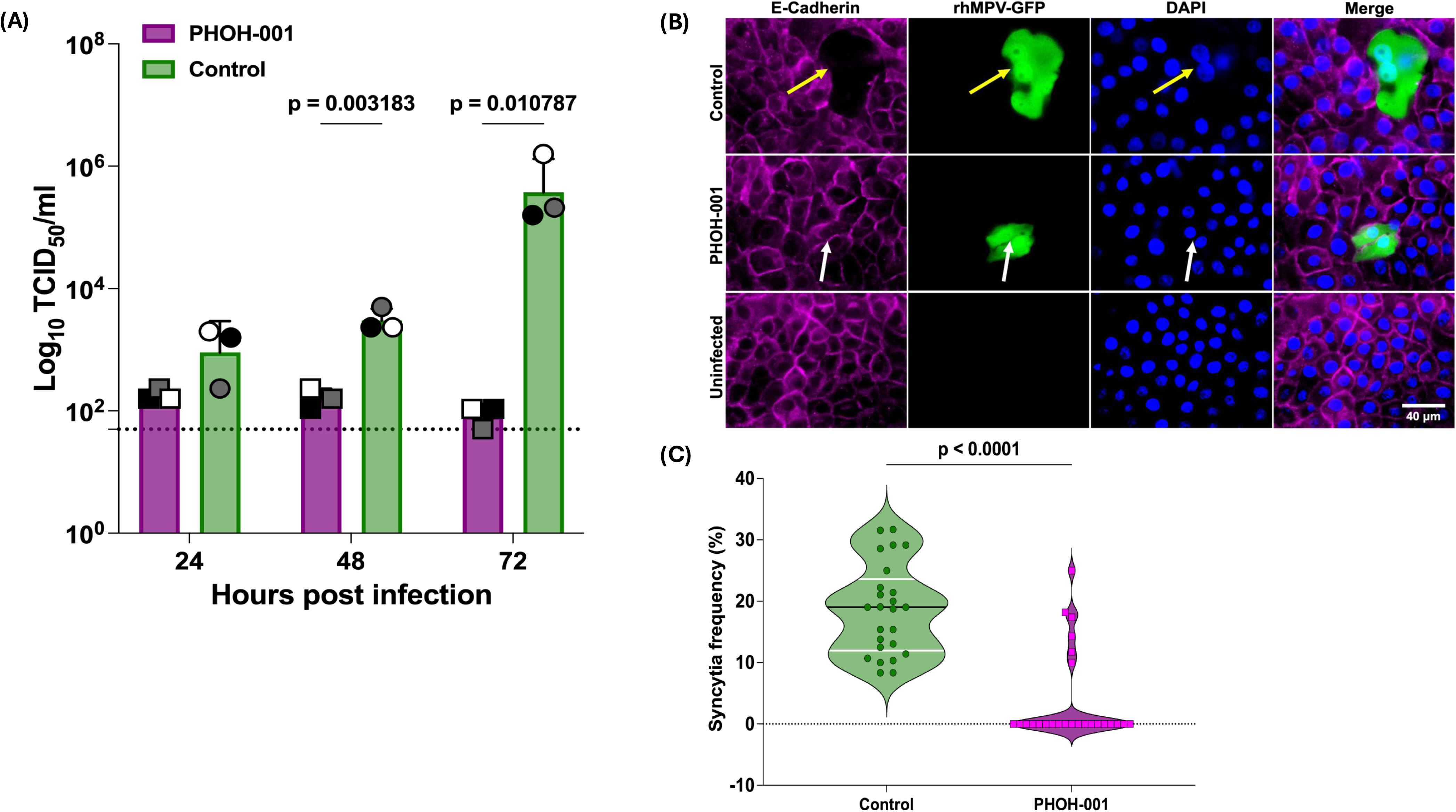
PHOH-001 significantly reduces hMPV replication and syncytium formation in HAECs. **(A)** TCID_50_ assay of hMPV replication in PHOH-001-treated HAECs. Infectious viral titers in supernatants of PHOH-001-treated and PBS-treated HAECs were measured by TCID_50_ assay at 24, 48, and 72 hpi. Data represents mean ± SD from three biological replicates, each with four technical replicates. Statistical significance between PHOH-001 and PBS at each time point was determined by multiple unpaired two-tailed t-tests with Bonferroni correction for multiple comparisons in GraphPad Prism (adjusted p<0.017). The dashed line indicates the LOD of the TCID_50_ assay (1.7 log_10_ TCID_50_/mL), and values below the LOD are plotted at the LOD. **(B)** Representative fluorescence images of infection models treated with or without PHOH-001 to analyze syncytia formation. HAECs were infected with rhMPV-GFP (MOI=1) and imaged at 60x magnification; syncytia are indicated by yellow arrows, and delineated (separate) infected cells are indicated by white arrows. Scale bar=40 µm. **(C)** Quantification of syncytium formation in HAECs. The percentage of DAPI-positive nuclei involved in syncytia (defined by GFP-positive areas, segmented based on E-cadherin, containing multiple nuclei) was quantified using fluorescence microscopy images and analyzed using CellProfiler software. Data are presented as violin plots, where individual points represent the percentage of total DAPI-positive nuclei that are part of syncytia; black line=median, white lines=upper and lower quartiles. Data represents mean ± SD from 25 technical replicates (p<0.05, two-tailed unpaired Student’s t-test).

### PHOH-001 alters hMPV F protein localization, limiting viral egress

The hMPV F protein is essential for viral entry and cell-to-cell spread. To assess whether PHOH-001 affects F protein trafficking, primary HAECs were infected with rhMPV-GFP (MOI=1) and treated with PHOH-001 or PBS. Low-magnification (20x) imaging revealed differences in actin organization between treatment groups, with PHOH-001-treated cells exhibiting altered cortical actin distribution compared to PBS controls (Figure 6A), consistent with previously reported actin reorganization during hMPV infection and studies demonstrating that actin dynamics influence pneumovirus replication and cellular architecture (40, 41). High-resolution confocal imaging at 72 hpi revealed marked differences in F protein distribution (Figure 6B). In PBS-treated cells, F protein was broadly distributed throughout the cytoplasm, consistent with normal intracellular trafficking and syncytium formation. In contrast, PHOH-001-treated cells exhibited pronounced enrichment of F protein at the cell periphery, proximal to the plasma membrane. This redistribution coincided with changes in cortical actin organization. Quantitative image analysis was performed using CellProfiler to assess F protein localization within GFP-positive infected cells. Phalloidin staining was used to delineate individual cell boundaries and define whole-cell and plasma membrane-proximal peripheral regions of interest, minimizing over-segmentation of multinucleated syncytia. Integrated F protein fluorescence intensity was measured on a per-cell basis, and peripheral enrichment was calculated as the ratio of peripheral to whole-cell intensity. PHOH-001 treatment significantly increased peripheral F protein enrichment relative to PBS controls (Figure 6C), indicating altered F protein trafficking and/or retention at the plasma membrane. Consistent with these localization changes, PHOH-001 reduced F protein expression at both the transcript and protein levels, as measured by RT-qPCR and western blot analysis (Supplemental Figure 4). Collectively, these findings suggest that PHOH-001 is associated with altered F protein localization and actin organization, consistent with impaired syncytium formation and reduced viral spread.

**Figure 6.**
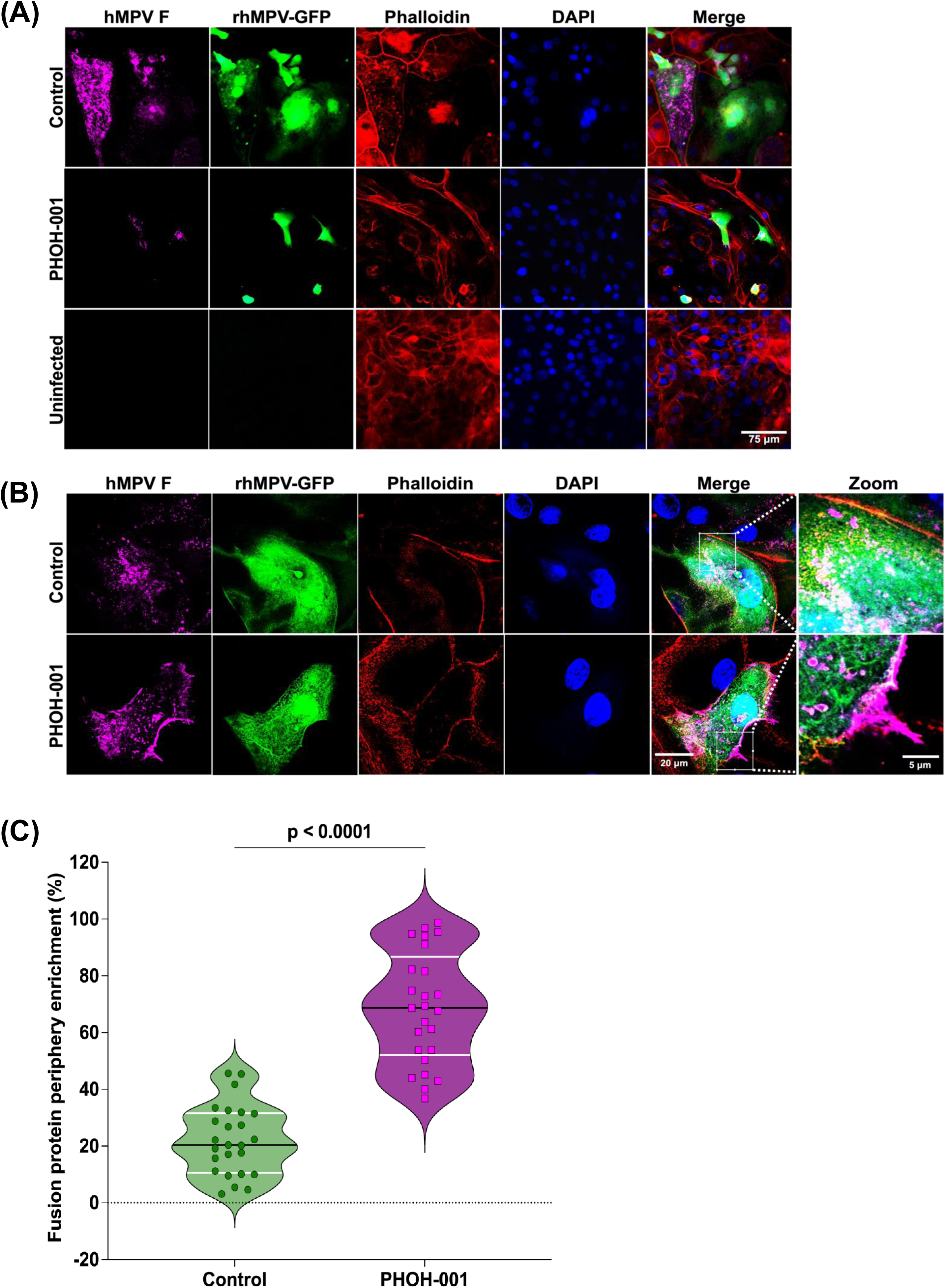
PHOH-001 alters hMPV F-protein localization. **(A)** Representative low-magnification (20x) images of primary HAECs infected with rhMPV-GFP (MOI=1) and treated with PHOH-001 or PBS at 72 hpi. Actin is visualized by phalloidin staining to delineate cellular morphology and was not independently quantified. PHOH-001 treatment is associated with altered actin organization compared to PBS controls. Scale bar=75 μm. **(B)** Representative high-resolution confocal images (63x) showing hMPV F protein distribution in infected HAECs at 72 hpi. In PBS-treated cells, F protein is broadly distributed throughout the cytoplasm and syncytia, whereas PHOH-001-treated cells exhibit pronounced enrichment of F protein at the cell periphery, proximal to the plasma membrane. Scale bar=20 μm. **(C)** Quantification of F protein peripheral enrichment within GFP-positive infected cells using CellProfiler. Individual cells were segmented based on phalloidin staining, and integrated F protein fluorescence intensity was measured within whole-cell and plasma membrane-proximal peripheral regions. Peripheral enrichment was calculated as the ratio of peripheral to whole-cell integrated intensity on a per-cell basis. PHOH-001 treatment resulted in a significant increase in F protein peripheral enrichment compared to PBS-treated controls. Data represents individual data points with mean ± SD from 25 technical replicates per condition (p<0.05, two-tailed unpaired Student’s t-test).

## Discussion

hMPV is a significant respiratory pathogen, particularly in immunocompromised individuals, the elderly, and young children, and there are currently no approved targeted therapeutics. Here, we demonstrate that PHOH-001, an inhaled alkaline buffer, effectively inhibits hMPV infection in primary HAECs under both submerged and ALI culture conditions, supporting modulation of airway epithelial pH as a potential antiviral strategy.

Our findings support a mechanistic model in which PHOH-001 interferes with multiple pH-dependent stages of the hMPV infection cycle (Figure 7). In untreated cells, hMPV can enter via endocytosis, where acidic endosomal pH promotes conformational activation of the viral F protein, enabling membrane fusion, genome release, and viral replication. Subsequent actin-dependent trafficking of F protein to the plasma membrane facilitates syncytium formation and viral egress, promoting efficient cell-to-cell spread. In PHOH-001-treated cells, elevated extracellular and intracellular pH likely impairs endosomal acidification and alters downstream trafficking-associated processes. Confocal imaging revealed accumulation of F protein at the cell periphery with reduced cytoplasmic dispersion and limited syncytium formation, correlating with markedly reduced infectious viral titers. Confocal imaging further demonstrated alterations in the cortical actin network following PHOH-001 treatment, consistent with prior reports that hMPV remodels actin to support intracellular trafficking and replication (38, 39). High-resolution imaging showed that although F protein localized proximal to the plasma membrane in PHOH-001-treated cells, it failed to achieve the broad membrane distribution characteristic of syncytium-competent infection. Together, these data indicate that PHOH-001 is associated with altered F-protein localization and actin organization, consistent with impaired viral replication, syncytium formation, and egress.

**Figure 7.**
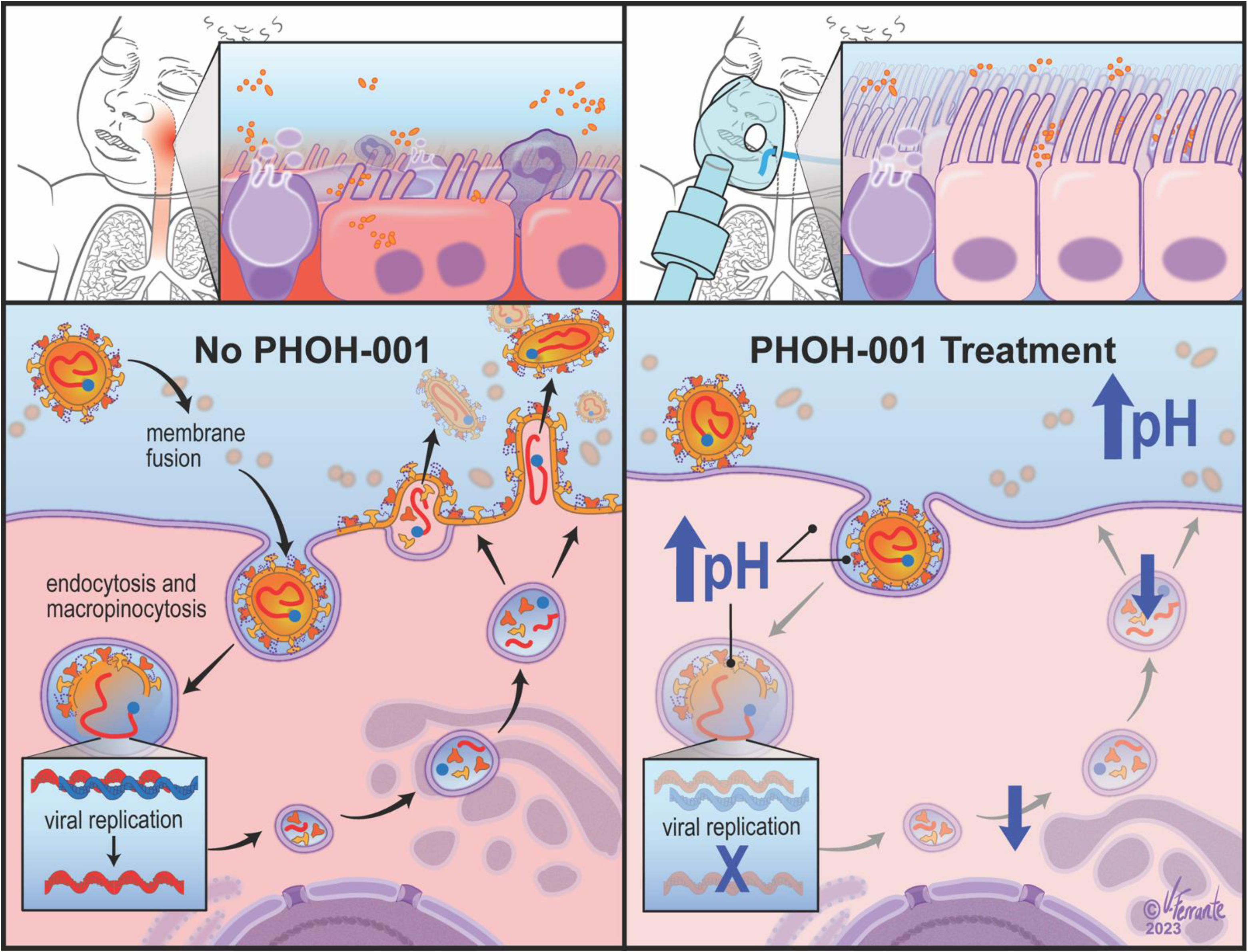
Proposed mechanism of PHOH-001’s action in modulating hMPV infection. Schematic representation of the hypothesized mechanism by which PHOH-001 modulates airway epithelial pH to inhibit hMPV infection. In the absence of PHOH-001 (left), hMPV can enter airway epithelial cells via endocytosis, where low pH in the endosome triggers conformational changes in the viral F protein, promoting membrane fusion and viral genome release. Subsequent F-protein trafficking to the plasma membrane, a process supported by the host actin cytoskeleton, facilitates syncytium formation and viral egress, enabling efficient viral spread. In the presence of PHOH-001 (right), elevated pH inhibits endosomal acidification, potentially reducing F-protein activation and viral fusion. In addition, altered actin organization may further disrupt F-protein trafficking and membrane dynamics, thereby limiting viral replication, syncytium formation, and egress, ultimately restricting viral spread and infection.

PHOH-001 has been evaluated *in vivo* and shown a favorable safety profile in both healthy and asthmatic individuals, with no reported adverse effects (33, 34). Consistent with these findings, prior studies in primary HAECs demonstrated that PHOH-001 does not induce cytotoxicity or disrupt epithelial integrity at concentrations used in antiviral experiments (33), supporting the interpretation that reduced hMPV infection observed here reflects antiviral activity rather than nonspecific cellular toxicity. Cytotoxicity was not directly assessed in the present study; however, treated HAECs retained normal morphology throughout the experimental period. Although *in vivo* efficacy against hMPV has not yet been demonstrated, the results from human safety trials support translational potential (33, 34). Furthermore, preliminary studies exploring PHOH-001 activity against other respiratory viruses, including SARS-CoV-2 and RSV, suggest that modulation of endosomal and airway pH may be a broadly applicable antiviral strategy (33, 35). Unlike direct antiviral inhibitors targeting viral proteins, modulation of airway pH represents a host-directed antiviral strategy that may reduce the likelihood of resistance development.

Limitations of this study include the use of a single hMPV A2 strain and exclusively *in vitro* HAEC models. Because pH sensitivity of hMPV fusion is strain dependent, evaluation across additional genotypes will be important to determine the breadth of this antiviral strategy. In addition, while reductions in syncytium formation and infectious viral titers suggest that PHOH-001 affects both early and post-entry stages of infection, the relative contribution of entry inhibition versus post-entry effects cannot be definitively resolved in the current study. Future studies incorporating time-of-addition approaches or direct measurements of entry and intracellular trafficking dynamics will be required to further define the stage(s) of inhibition.

In summary, PHOH-001 represents a novel approach to modulate airway epithelial pH and inhibit hMPV infection by altered F-protein localization, actin-associated trafficking, syncytium formation, and viral egress. Our findings provide a mechanistic foundation and strong rationale for advancing PHOH-001 toward preclinical and clinical evaluation as a potential therapeutic for hMPV and other pH-dependent respiratory viral infections.

## Acknowledgements

We gratefully acknowledge Dr. Sabrina Absalon for technical assistance with microscopy. We acknowledge the use of BioRender.com for figure schematics, and the software tools CellProfiler and Fiji for imaging analysis.

## Author Contributions

I.A.D. conceived the study, performed experiments, acquired and analyzed data, secured funding, and wrote the original draft and subsequent edits. A.L. contributed to methodology, data analysis, and manuscript editing. J.S. assisted with conceptualization, methodology, and data analysis. L.S. contributed to methodology and maintenance of cell cultures. N.T. assisted with methodology and manuscript editing. R.F.R. provided conceptual input and manuscript editing. T.E. contributed to maintenance of cell cultures. B.G. contributed to conceptualization, supervision, manuscript editing, and funding acquisition. M.D.D. provided conceptual guidance, supervision, and manuscript editing. All authors approve manuscript for publication.

## Supplemental Data

**Supplemental Table 1.** Donor demographics for primary HAECs used including age, sex, and health status.

**Supplemental Table 2.** Primer, probe, and gene block design for CAN97-83 [A2] fusion protein RT-qPCR.

**Supplemental Figure 1. Validation of rhMPV-GFP replication kinetics in Vero E6 cells. (A)** Representative fluorescence images of Vero E6 cells infected with rhMPV-GFP at MOI 0.01, 0.1, or 1. Images were captured at 18, 24, 48, and 72 hpi at 2x magnification. Scale bar=1250 µm. **(B)** Dual-axis quantification of GFP signal (RLU; left y-axis) and infectious titers (TCID_50_/mL; right y-axis) at 18, 24, 48, and 72 hpi. GFP signal was measured directly from the assay plate using a plate reader, and infectious titers were determined from supernatant collections by TCID_50_ assays on Vero E6 cells. Data represents mean ± SD from three technical replicates. RLU and TCID_50_ increased proportionally with MOI, validating assay performance and supporting subsequent application in primary HAECs. The dashed line indicates the LOD of the TCID_50_ assay (1.7 log_10_ TCID_50_/mL). Values below the limit of detection were plotted at the LOD.

**Supplemental Figure 2. Plate-based imaging of pHrodo™ Red fluorescence following PHOH-001 treatment. (A)** Representative whole-plate image showing pHrodo™ Red fluorescence in Vero E6 cells treated with PHOH-001 (120 mM) followed by 1:2 serial dilutions across a 96-well plate. Images were acquired with an Odyssey M Imager immediately following fluorescence quantification by plate reader to confirm consistency between fluorescence intensity and spatial signal distribution. The image shown corresponds to a single row of the 96-well plate from this experiment.

**Supplemental Figure 3. Western blot and RT-qPCR validation of dose-dependent hMPV inhibition by PHOH-001. (A)** Western blot analysis of primary HAECs infected with rhMPV-GFP (MOI=1) and treated with increasing concentrations of PHOH-001. A narrower dose range was used to validate the fluorescence-based inhibition observed in Figure 4. Blots show reduced GFP expression with increasing PHOH-001 concentration. **(B)** Quantification of GFP expression normalized to β-actin confirms a significant, dose-dependent reduction in GFP levels with PHOH-001 treatment. Data represents four technical replicates and were analyzed using one-way ANOVA with post hoc Tukey’s test (p<0.05). **(C)** RT-qPCR assay performance was validated using a standard curve generated from serial dilutions of the hMPV F protein gene amplicon. **(D)** RT-qPCR quantification of the hMPV F protein gene revealed a clear dose-dependent antiviral effect of PHOH-001. As PHOH-001 concentration increased, F protein gene expression progressively decreased, indicating strong inhibition of hMPV replication at higher PHOH-001 doses. The RT-qPCR is representative of three technical replicates and data were analyzed using one-way ANOVA with post hoc Tukey’s test (p<0.05).

**Supplemental Figure 4. Western blot and RT-qPCR confirmation of reduced F gene and protein expression in HAECs treated with PHOH-001. (A)** Western blot analysis of primary HAECs infected with rhMPV-GFP (MOI=1) comparing PHOH-001–treated and PBS-treated groups. Blots show reduced hMPV F protein levels in PHOH-001–treated samples relative to controls. **(B)** Quantification of hMPV F protein expression normalized to β-actin confirms a significant reduction in PHOH-001–treated samples. Data represents three technical replicates and were analyzed using one-way ANOVA with post hoc Tukey’s test (p<0.05). **(C)** Standard curve generated from serial dilutions of the hMPV F gene amplicon to validate RT-qPCR assay performance. **(D)** RT-qPCR analysis of hMPV F gene expression in PHOH-001–treated versus untreated HAECs. Results show a significant decrease in fusion gene transcript levels in the treatment group, consistent with the protein-level reduction observed in panel A. Data represents six technical replicates and were analyzed using one-way ANOVA with post hoc Tukey’s test (p<0.05).

