## Supplemental Data for "A novel antiviral strategy targeting human metapneumovirus through pH modulation in human airway epithelial cells"

#### Supplemental Table 1

| Donor # | Sex, Age | Disease State | Origin |
|---|---|---|---|
| #07783 | M, 12 YO | Healthy | Bronchial |
| #0059 | F, 25 YO | Healthy | Bronchial |
| #0098 | M, 42 YO | Healthy | Bronchial |

#### Supplemental Table 2

| Name: | Sequence(5'- 3') |
|---|---|
| Forward Primer (fusion protein) | AATTGCCAAAACCATCCGGC |
| Reverse Primer (fusion protein) | CACTGCAGTTGCCAACACTC |
| Probe (fusion protein) | 6FAM-TCACAGCAATTAAGAATGCCCTCAAACGACCAATGAA-MGBNFQ |
| Gene Block (fusion protein) | AATAGCACTCGGTGTTGCAACAGCAGCTGCAGTCACAGCAGGTGTTGCAATTGCCAAAACCATCCGGCTTGAGAGTGAAGTCACAGCAATTAAGAATGCCCTCAAAACGACCAATGAAGCAGTATCTACATTGGGGAATGGAGTTCGAGTGTTGGCAACTGCAGTGAGAGAGCTGAAAGACTTTGTGAGC |

#### Supplemental Figure 1

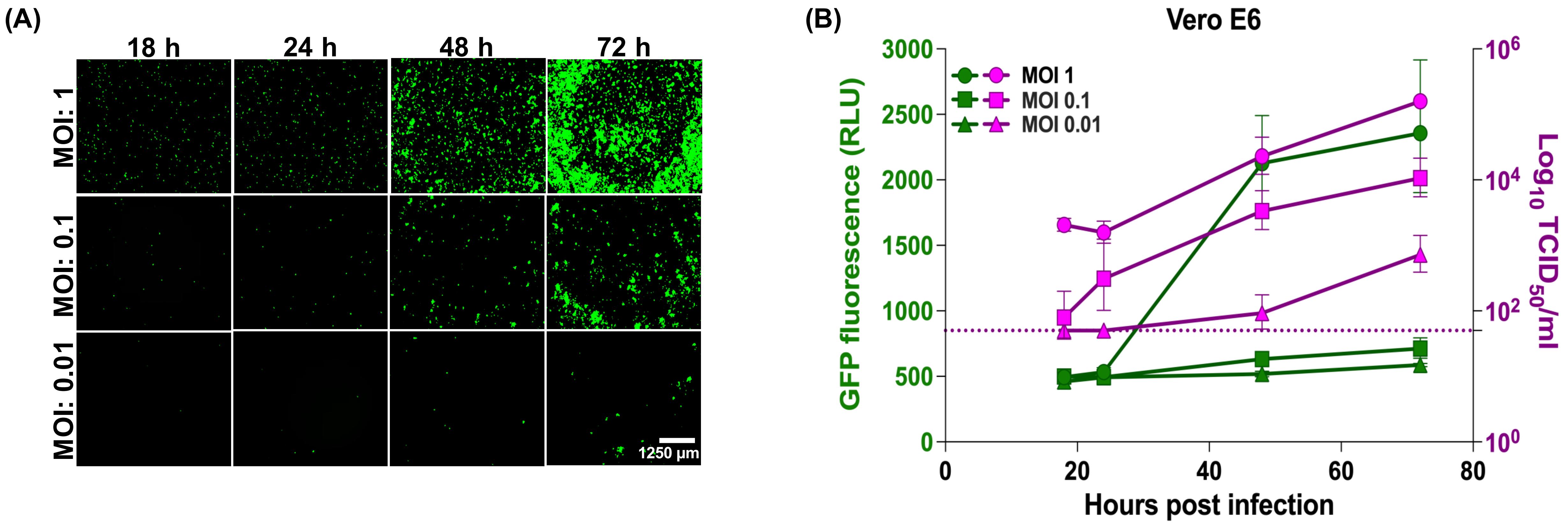

### Supplemental Figure 2

(A)

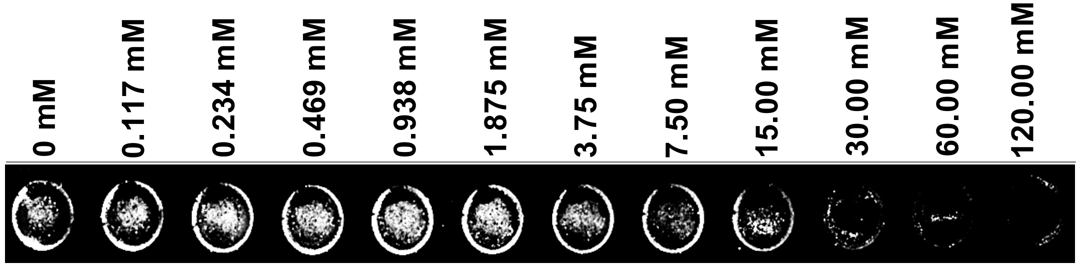

### Supplemental Figure 3

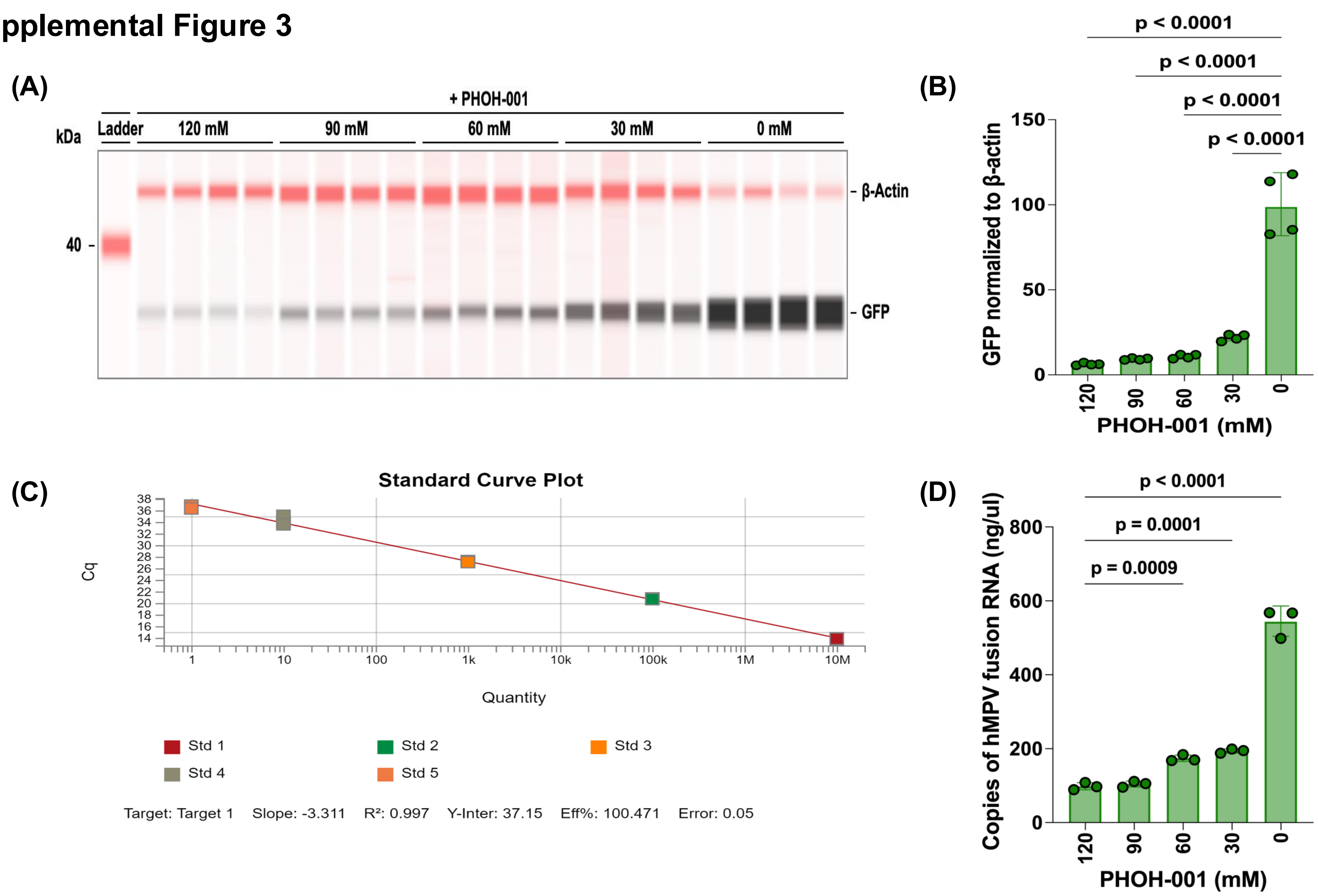

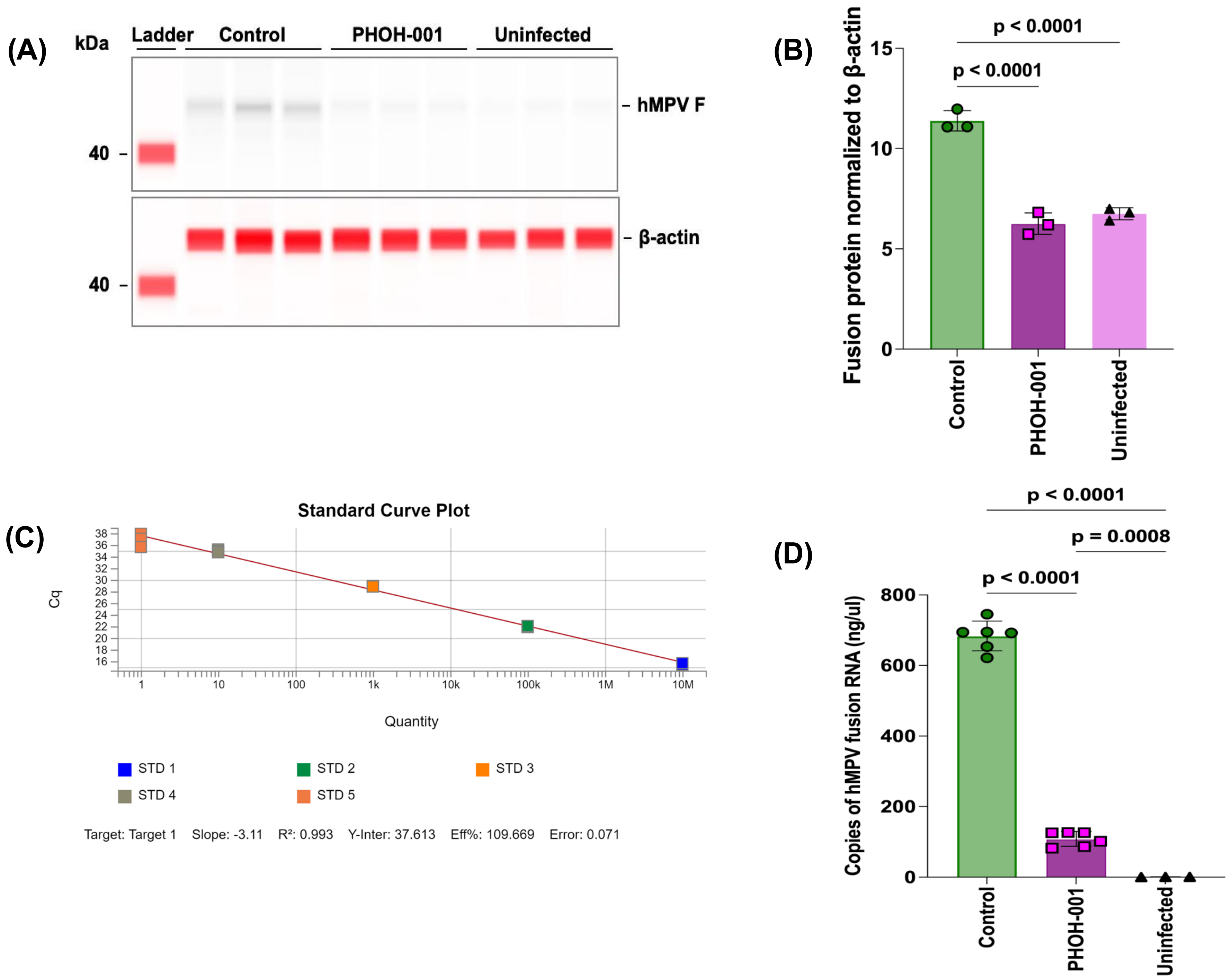


Supplemental Figure 4
